# DNA-free CRISPR Genome Editing in Raspberry (*Rubus idaeus*) through RNP-mediated Protoplast Transfection and Comparison of Indel Analysis Techniques

**DOI:** 10.1101/2025.01.14.632935

**Authors:** Ryan Creeth, Andrew Thompson, Zoltan Kevei

## Abstract

Protoplast-based systems have been utilised in a wide variety of plant species to enable genome editing without chromosomal introgression of foreign DNA into plant genomes. This allows elite cultivars to be edited without further genetic segregation, preserving their unique genetic composition and their regulatory status as non-transgenic. This can be achieved by DNA-free genome editing in protoplasts, followed by regeneration. However, protoplast isolation presents a barrier to the development of advanced breeding technologies in raspberry and no protocol has been published for DNA-free genome editing in the species. Pre-assembled ribonucleoprotein complexes (RNPs) do not require cellular processing and the commercial availability of Cas9 proteins and synthetic guide RNAs has streamlined genome editing protocols. This study presents a novel high-yielding protoplast isolation protocol from raspberry stem cultures and RNP-mediated transfection of protoplast with CRISPR-Cas9. Targeted mutagenesis of the phytoene desaturase gene at two intragenic loci resulted in an editing efficiency of 19%, though estimated efficiency varied depending on the indel analysis technique. Only amplicon sequencing was sensitive enough to confirm genome editing in a low efficiency sample. To our knowledge, this study constitutes the first use of DNA-free genome editing in raspberry. This protocol provides a valuable platform for understanding gene function and facilitates the development of precision breeding in this important soft fruit crop.

## Introduction

Raspberry (*Rubus idaeus*) is a high-value horticultural crop that, alongside other *Rubus* berries, has undergone a substantial increase in consumption worldwide in recent decades ^1^. Globally, production increased 48% between 2011 and 2021 ^2^. Often sold as a ‘superfood’, raspberries contain high levels of essential vitamins and bioactive compounds including anthocyanins, flavanols and phenolic acids that are essential for normal metabolism. Lack of dietary sources of these bioactive nutrients has been associated with cancer, stroke, Alzheimer’s and autoimmune diseases ^3,4^. Raspberry is a highly heterozygous species that is not true to seed, meaning both sexual reproduction and selfing can introduce allelic variation that alters the berry phenotype of elite cultivars. Commercial raspberry production thus relies on vegetative propagation. This challenge, in addition to a two-year fruiting cycle in floricanes, has contributed to a lack of progress on next-generation methods for raspberry breeding.

Improving agricultural sustainability is a key priority for reaching net zero ^5^. Food production systems must undergo sustainable intensification, and gains can be made in raspberry production through improving efficiency while reducing food waste. Advanced breeding to improve traits such as fungal resistance or fruit firmness would substantially improve shelf life ^6^. Enhanced plant and fruiting architecture could reduce labour demands for crop management and harvesting; eliminating genetic disorders, such as crumbly fruit syndrome, would prevent widespread yield losses ^7^. Such improvements can also minimise chemical inputs and decrease overall land use, leading to more productive and sustainable raspberry cultivation.

The advent of CRISPR (clustered regularly interspaced short palindromic repeats) genome editing has greatly increased the scope and speed of plant breeding ^8^. Through the formation of ribonucleoprotein complexes (RNPs), composed of Cas nucleases and programmable guide RNA (gRNA), targeted mutagenesis of any genomic region of interest can be achieved ^9–11^. The most commonly used nuclease, Cas9, induces a double strand break (DSB) in genomic DNA at a specific locus identified by complementary binding of the gRNA ^12^. Often, error-prone DNA repair subsequently creates insertions and/or deletions (indels) at the specified locus, leading to gene knockout.

Applying this system to plant species is challenging as the plant cell wall is impermeable to nucleic acids encoding Cas9/gRNA or pre-assembled RNPs. Inbreeding, annual, seed-propagated crops can utilize *Agrobacterium*-mediated transformation or biolistics to transfer DNA encoding CRISPR/Cas9 into intact cells and to integrate this into chromosomes. After genome editing has occurred, transgenes can be easily removed by backcrossing and genetic segregation as transgenic crops have historically been contentious from regulatory and public perception standpoints ^13,14^. However, to realise the benefits of genome editing in the advanced breeding of allogamous, clonally-propagated crops, such as raspberry, editing must take place without chromosomal introgression of foreign DNA. This allows elite clones to be edited without the need for further genetic segregation, preserving their unique genetic composition and their regulatory status as non-transgenic. Where *Agrobacterium*-mediated transformation cannot be used, the resurgence of protoplast-based techniques offers a transgene-free alternative ^15–20^.

Enzymatic digestion of the plant cell wall (enzymolysis) liberates single-celled protoplasts ^16,21,22^ with membranes permeable to plasmids and pre-assembled RNPs in the presence of polyethylene glycol (PEG) ^20^. After editing, protoplast can be regenerated into whole plants ^18,23,24^, although this is technically challenging and species dependent. Cas9 and gRNA genes encoded within plasmids are expressed within the protoplast nucleus, triggering genome editing, but do not integrate into the genome. Pre-assembled RNPs, exogenously synthesised and formed extracellularly, are also able to directly translocate the protoplast membrane to initiate editing ^20^. RNPs are then degraded by normal cellular processes. This form of DNA-free genome editing is rapid, highly specific and indistinguishable from natural mutagenesis – factors that have recently found favour with regulators ^25,26^. Pre-assembled RNPs are of particular interest in crop species like raspberry as they eliminate the chance of random integration of DNA into the genome, which is a risk with plasmid transfection. The commercial availability of synthetic gRNAs and Cas nucleases from many vendors, and evidence for higher editing efficiencies than plasmid-based techniques ^27^, also favour the use of DNA-free RNPs.

Genome editing and genetic engineering in raspberry have not been extensively developed ^1^. *Agrobacterium*-mediated transformation has been achieved by several groups ^28–32^, however transgenic raspberries have never been commercialised. To our knowledge, no method for DNA-free CRISPR genome editing in raspberry has been published, however there have been attempts to use CRISPR transgenically ^32,33^, with some potential success with biolistic delivery. Nonetheless, there is a substantial research gap in this field. Pre-assembled RNPs present an excellent opportunity to rapidly test gRNAs if reliable and modern protoplast-based methods can be elucidated. Existing protoplast research on raspberry is limited in scope and predates the advent of genome editing ^34–36^, but provides evidence that raspberry protoplasts can be isolated and cultured. Many raspberry cultivars are suitable for tissue culture; vegetative propagation is common industry practice ^37^ and raspberry produces shoots vigorously, including directly from roots, suggesting a propensity for regeneration.

This study demonstrates successful, DNA-free genome editing of raspberry protoplasts using CRISPR-Cas9 for the first time. We have applied state-of-the-art protoplast methods to a new species and achieved a high genome editing efficiency. These are critical steps in the development of next generation breeding technologies (NGBTs) in raspberry and will hopefully stimulate further work in an underserved yet economically and nutritionally valuable species.

## Methods

### Plant Cultivation

Cold-treated (4°C, ≥50 days, ∼15×20 cm) roots of *R. idaeus* cultivar BWP102 (supplied by a commercial propagator) were planted in 7.5 L pots containing peat-rich soil (Sinclair All Purpose Growing Medium) with perlite and fertilised with Hoagland’s solution ^38^ twice a week. Highly vigorous, 1.5 cm diameter, 1 m tall canes produced from cold-treated roots were cut down after approximately one month in a partially environmentally controlled glasshouse (set points of 20°C, 16:8hrs day: night, supplemented with high-pressure sodium lamps). Petioles and leaves were removed, and cane stems cut into 50-70 mm segments containing one axillary bud. Cuttings were sterilised in 70% v/v ethanol for 30 seconds and 10% v/v sodium hypochlorite (with 0.002% Tween 20) for four minutes ^37^. Exposed stem ends were cut off and stem explants were planted in Murashige and Skoog (MS) basal media with Gamborg B5 vitamins (Melford; pH 5.8, 30 g L^-1^ sucrose, 7.5 g L^-1^ agar, 1 mg L^-1^ 6-benzyl-aminopurine, 0.1 mg L^-1^ indole-3-butyric acid) within autoclaved Magenta™ boxes (Merck).

Tissue culture material was grown in a Sanyo MLR-350 growth cabinet with 45 µmol m^-2^ s ^-1^ light from fluorescent tubes (Toshiba FL40SSW/37) at 25°C with a 16: 8hr day: night cycle.

### Protoplast Isolation

Protoplasts were isolated from plantlets (stem cultures) grown from the axial bud cane cuttings. Plantlets (∼0.3 g, 20-30 mm tall) from axillary buds were harvested after 10 days of culture and cut with a sterile scalpel blade into 0.5-1 mm strips, then immediately immersed in 13 mL enzymolysis solution (20 mM MES pH 5.8, 0.4 M mannitol, 20 mM KCl, 10 mM CaCl_2_, 1.5% Cellulase R-10 (Duchefa), 0.5% Macerozyme R-10 (Duchefa)) in a 90 mm Petri dish^16^. The tissue was vacuum infiltrated for 30 minutes and then incubated on an orbital shaker at 50 rpm for 16 hrs in the dark at 28°C. Protoplast were harvested by adding 10 mL of W5 media (2 mM MES pH 5.8, 154 mM NaCl, 125 mM CaCl_2_, 5 mM KCl) to the Petri dish and agitating gently, then filtering the suspension through a 70 µM cellulose filter (Fisher Scientific) into a 50 mL centrifuge tube. Protoplast were pelleted by centrifugation (Sigma 3-16KL) at 200 rcf for four minutes with slow centrifugal acceleration and breaking (both set at level 6). The supernatant was removed, and protoplasts were resuspended in W5 for a second wash at 200 rcf for four minutes. Finally, the supernatant was removed, and 2 mL 0.4 M mannitol (pH 5.8) was added to resuspend the protoplast.

Protoplast were then purified through a sucrose cushion ^23^: 6 mL of 0.6 M sucrose solution was gently overlaid with the 2 mL protoplast-mannitol suspension in a 15 mL centrifuge tube and centrifuged at 80 rcf for five minutes. Protoplast suspended at the interface were carefully aspirated into a new 15 mL tube and then centrifuged at 200 rcf for four minutes and resuspended in 5 mL MMG (4 mM MES pH 5.8, 0.4 M mannitol, 15 mM MgCl_2_) solution to form a highly concentrated pellet of viable protoplast. Sucrose purification was only successful when initial protoplast yields exceeded 1 × 10^6^ protoplast mL^-1^.

### *In vitro* Cleavage Test and gRNA Design

PCR product of BWP102 *phytoene desaturase* (*PDS*) was used to test the efficacy of two synthetic gRNAs that targeted separate regions of the gene *in vitro. PDS* in raspberry was identified by BLAST mapping strawberry (*Fragaria vesca*) *PDS* sequence to a novel genome assembly of raspberry cultivar BWP102 (Kevei et al., unpub.) in Geneious Prime v.2024.0.7 (Dotmatics). gRNAs and primers were designed using the same genome assembly to amplify two separate coding regions of *PDS* (*PDS1, PDS2*) with gRNA cut sites producing uneven cleavage products distinguishable by gel electrophoresis (Table 1). gRNA1 and gRNA2 targeted separate loci in exon 13 and eight of *PDS* respectively. Custom Invitrogen TrueGuide gRNAs and Invitrogen TrueCut Cas9 were commercially synthesised by Thermo Fisher Scientific.

**Table 1:**
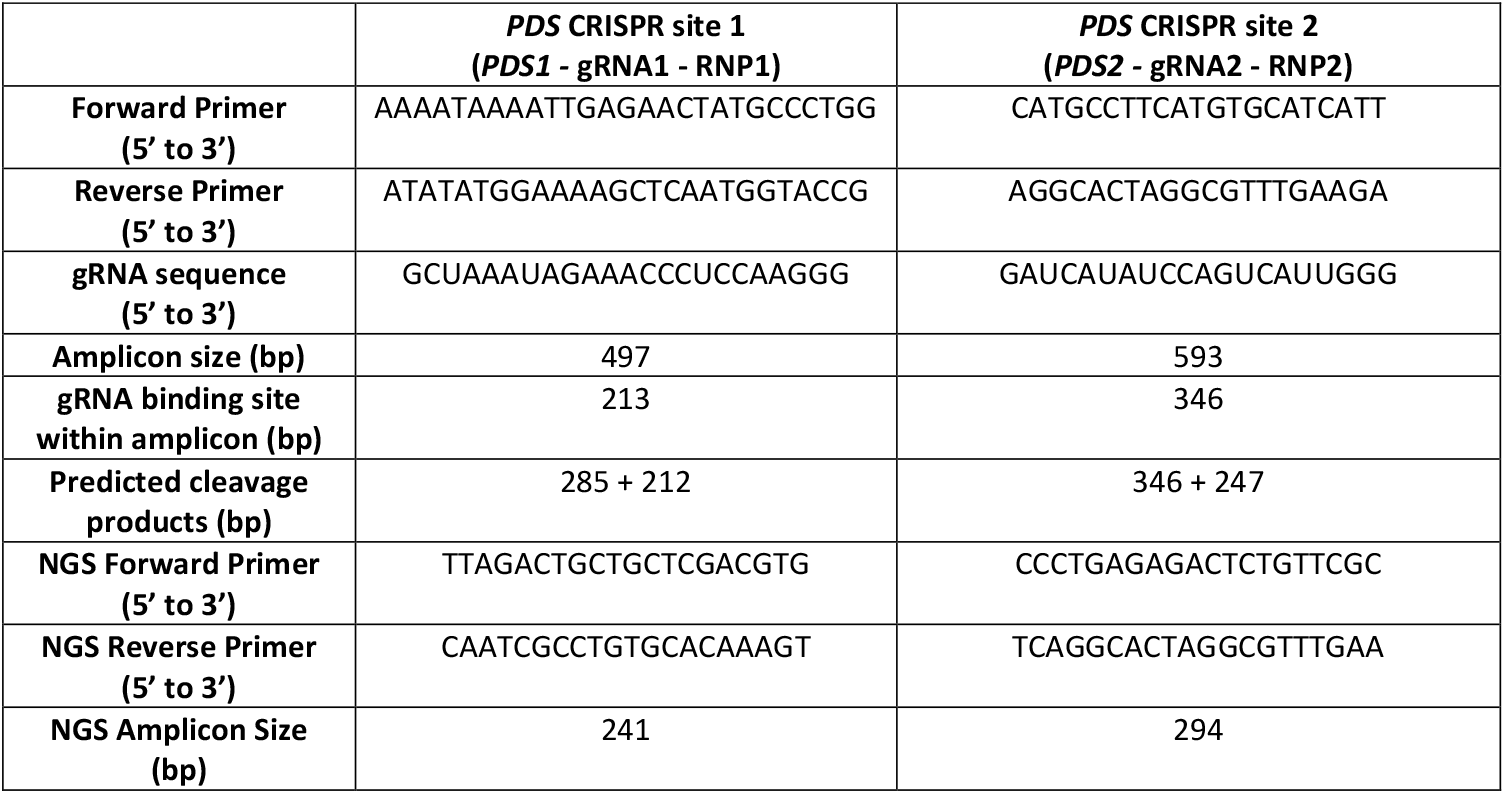
Oligonucleotides and gRNAs used in this study. Predicted cleavage products were deduced assuming DSB 3bp upstream from PAM site. NGS primer pairs flank the same respective gRNA binding site. *PDS*, phytoene desaturase; gRNA, guide RNA; RNP, ribonucleoprotein complex; bp, basepair; NGS, next generation sequencing.

DNA was extracted from BWP102 leaf tissue with E.Z.N.A Plant DNA kit (Omega Bio-tek) and *PDS* amplified with Phusion high-fidelity polymerase (Thermo Fisher Scientific), followed by purification with QIAquick PCR purification kit (QIAGEN). The *in vitro* cleavage test was followed as in Brandt et al. (2020) ^39^. In brief, 1 µg gRNA and 1 µg Cas9 were pre-mixed and incubated with 200 ng *PDS* PCR product for two hours. The sample was run on a 2% agarose gel (0.5% Tris Borate EDTA buffer, 0.005% Safeview (NBS Biologicals)). RNP activity was identified by the cleavage of the PCR product into two distinct smaller bands of expected sizes.

### Protoplast Transfection

The transfection protocol was based on Park et al. (2019) ^40^ with modifications. The protoplast suspension was centrifuged at 200 rcf for four minutes with low acceleration and deceleration after sucrose cushion filtration. The supernatant was removed as much as possible; 0.05% fluorescein diacetate (FDA) was added to a 10 µL subsample and protoplast yield was determined via haemocytometry on a fluorescent microscope (Leica Microsystems, DM2500). Protoplast concentration was standardised to 1 × 10^6^ cells mL^-1^ in MMG and 200 µL (2 × 10^5^ protoplast total) was transferred to a sterile 1.5 mL microcentrifuge tube using wide-bore pipette tips.

RNPs were formed by mixing 10 µg of gRNA, 10 µg Cas9, 2 µL lipofectamine CRISPRMAX (Thermo Fisher Scientific), 2 µL Plus reagent, 2 µL NEB 3.1 buffer (New England Biolabs) up to 20 µL with ultrapure H_2_O in a PCR tube and incubated for 10 minutes at 25°C. Negative controls were identical but excluded gRNA. RNPs were added to the protoplast suspension followed immediately by 220 µL 40% PEG4000 (Merck) solution (0.2 M mannitol, 0.1 M CaCl_2_, 2 g PEG4000, up to 5 mL ddH_2_O). The RNP-protoplast suspension was pipetted 5-10 times with a wide-bore tip and incubated for 10 minutes at 25°C. Transfection was terminated by the addition of 800 µL of W5 and the suspension transferred to a clean 15 mL tube and centrifuged for two minutes at 200 rcf. The supernatant was discarded and 800 µL W5 added for a second wash through centrifugation for two minutes at 200 rcf. The supernatant was removed and 500 µL of basic culture media (MS pH 5.8, 0.4 M mannitol, 30 g L^-1^ sucrose) was added to resuspend the protoplast and the suspension was left for 24 hrs. Protoplast were lysed by centrifugation at 10,000 rcf for one minute, the culture media was aspirated, and DNA was then extracted with the E.Z.N.A Plant DNA kit (Omega Bio-tek).

### Determination of Editing Efficiency with T7EI and Deconvolution

Editing efficiency was determined both *in vitro* and *in silico*. Protoplast DNA was amplified by PCR with Phusion polymerase with the same respective primer pairs used for the *in vitro* cleavage test. Amplification was visualised on a 2% agarose gel. PCR products were purified with QIAquick PCR purification kit and the T7 endonuclease I (T7EI) digestion protocol was followed ^41^ with minor modifications. In brief, PCR amplicons were denatured at 98°C for 30 seconds and then annealed by decreasing the temperature gradually by −2°C s^-1^ to 85°C, then −0.1°C s^-1^ to 25°C. T7EI (New England Biolabs) was added and incubated for one hour at 37°C ^39^. Annealed amplicons were purified again and then run on a 2% agarose gel for 45 minutes. PCR product from the same genome edited protoplast DNA samples was purified and sent to GENEWIZ (Azenta Life Sciences) for Sanger sequencing. Files from negative control and experimental samples were uploaded onto Tracking of Indels by Decomposition (TIDE) and deconvoluted ^42^ to estimate editing efficiency.

### Determination of Editing Efficiency with NGS

PCR product from the same genome edited protoplast DNA samples described for T7EI/deconvolution was purified and also sent to GENEWIZ (Azenta Life Sciences) for short-read Illumina next generation sequencing (NGS; Amplicon-EZ). Different primer pairs flanking the same gRNA binding sites were used for NGS as Amplicon-EZ requires primer pairs <450 bp (Table 1). FASTQ. files from the amplicon sequencing (≤70k sequences) were paired, trimmed and merged. Total wild-type and variant sequence percentages were compared to identify CRISPR indels in Geneious Prime ^43^.

## Results

### Tissue Culture and Protoplast Isolation

The quality of the initial raspberry canes was a critical factor in protoplast isolation (Figure 1a, b). Cuttings from highly vigorous, one-month old canes with no signs of senescence (no wilting/discolouring of any leaves, deep red thorns, thick bright green stem) generated fast growing plantlets (Figure 1c) that yielded between 1 × 10^6^ to 1.2 × 10^7^ protoplasts ml^-1^, with an average of ∼5 × 10^6^ protoplast ml^-1^ (Figure 1f, g).

**Figure 1:**
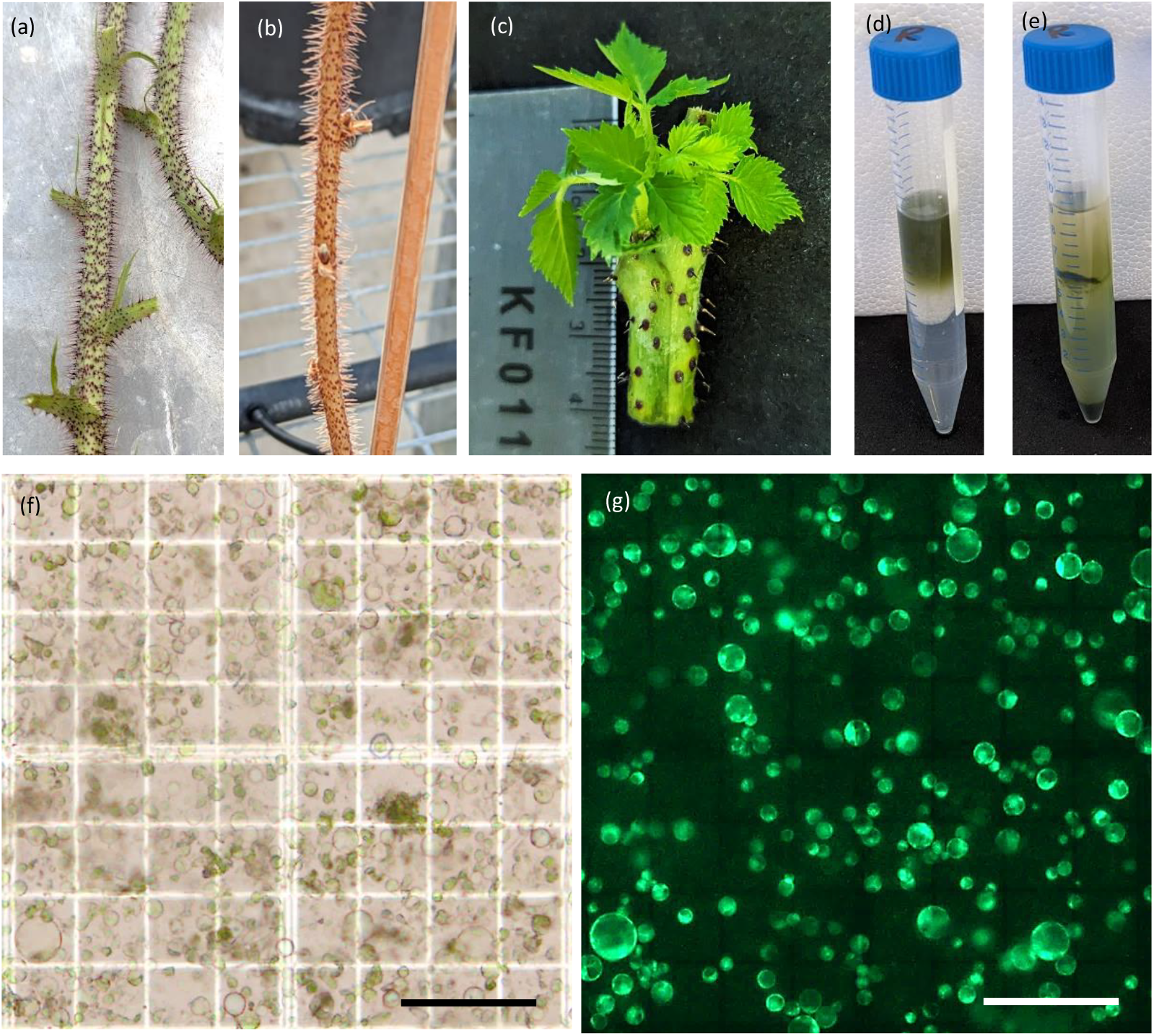
Protocol for protoplast isolation in raspberry. (a) High-quality canes used for shoot culture, (b) low-quality cane, (c) young plantlets formed from shoot cultures, (d, e) purification of protoplast before (d) and after (e) sucrose cushion, (f, g) protoplast isolated from plantlets under bright-field (f) and fluorescent (g) illumination (bars = 100µM).

Protoplast isolated from plantlets were 10-40 µM in diameter and bright green in colour. The sucrose cushion allowed the concentration of viable protoplast and removal of debris and dead cells. FDA staining demonstrated that the vast majority of protoplast were viable for transfection.

### *In vitro* Cleavage of DNA Amplicons by RNPs

Both RNPs showed a high level of activity against PCR amplicons with two bands clearly visible below the original band (Figure 2a, b). Compared to negative controls, the original PCR product was much reduced in intensity – indicating almost complete cleavage of the amplicons. RNP1 and RNP2 cleavage resulted in predicted bands at approximately 212 bp + 285 bp and 247 bp + 346 bp, respectively. Cleavage by the RNPs was highly specific as no other bands were visible (Figure 2a, b).

**Figure 2:**
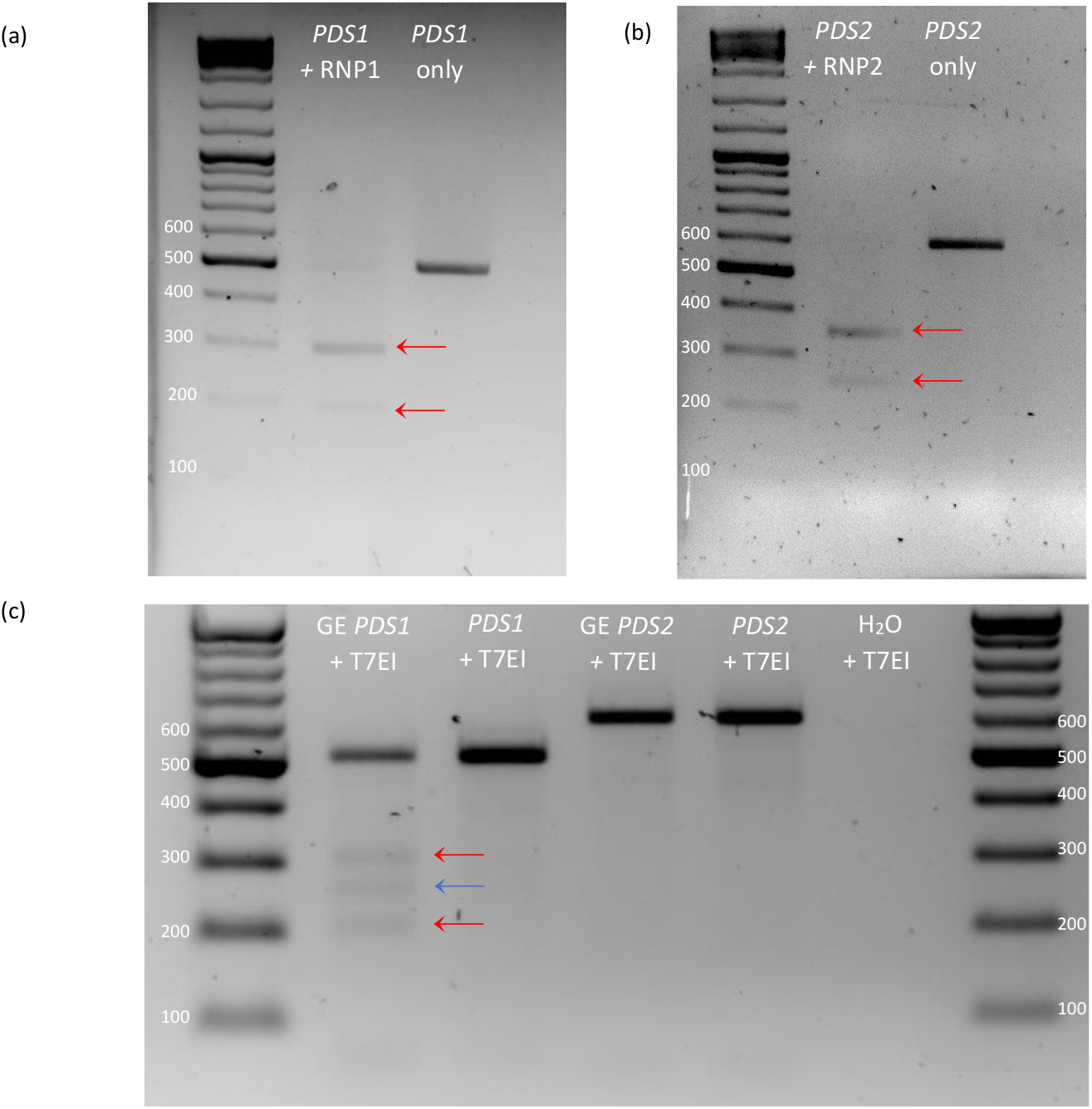
Detection of *in vitro* RNP activity and genome editing in protoplasts. (a, b) *In-vitro* cleavage test of RNP1 (a) and RNP2 (b) on leaf-derived *PDS* PCR product. (c) T7EI assay of *PDS* PCR product of gene edited (GE) protoplast transfected with RNP1 and RNP2. Red arrows indicate predicted cleavage products, and the blue line indicates the additional cleavage product.

### Detection of Genome Editing in Protoplasts

The T7EI assay confirmed the presence of mutations in the *PDS1* sample through the presence of cleavage products beneath the WT amplicon band. Two bands were approximately the predicted sizes at 212 bp and 285 bp, however there was an additional band at 250 bp. Furthermore, the WT amplicon band was also higher than expected at approximately 510 bp. However, *PDS2* showed no cleavage by T7EI (Figure 2c).

TIDE deconvolution estimated editing efficiencies of 14% for *PDS1* with predicted indels ranging from +1 to −5 (Figure 3a). Despite the T7EI assay not producing any visible cleavage bands, editing efficiency was estimated at 6.2% for *PDS2* with the most common predicted indel of +1 (Figure 3c). Aberrant signal increased directly downstream of the gRNA binding site, particularly for *PDS1*, which was also visible on the chromatogram (Figure S1a, c).

**Figure 3:**
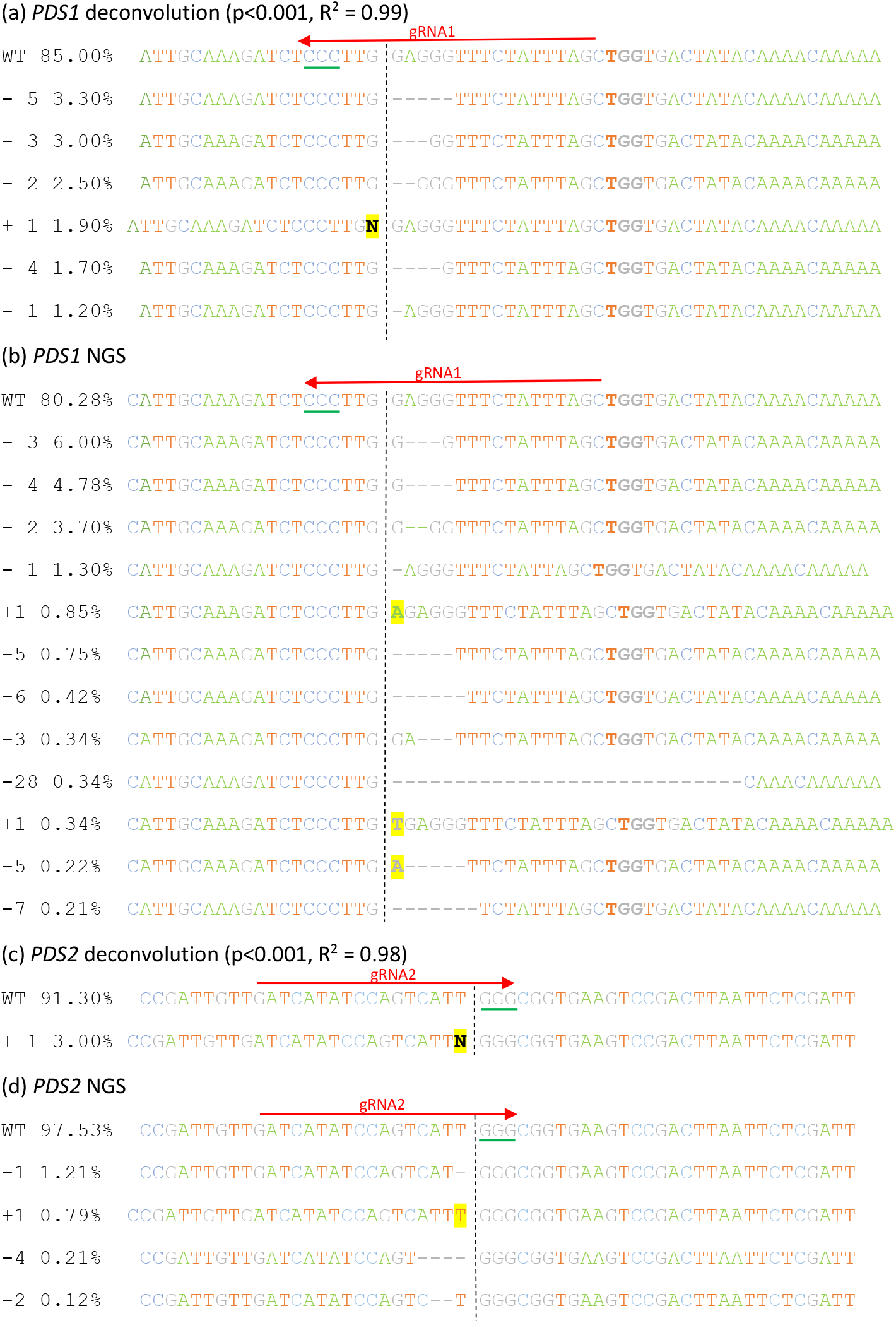
Estimated indels predicted by Sanger sequencing/ TIDE deconvolution and actual indels detected by NGS. (a, b) *PDS1* indels (c, d) *PDS2* indels. N refers to estimated insertions of unknown nucleotides from TIDE analysis. Vertical dashed lines indicate gRNA cut sites, red arrows indicate gRNA sequences and green lines indicate PAM sequences. All sequences are shown in the 5’ to 3’ direction. Note that gRNA1 binds in the 3’ to 5’ direction.

Amplicon sequencing followed by CRISPR analysis revealed 19.0% editing efficiency at *PDS1* with 12 variants (minimum presence in sequence population of 0.1%) ranging from −28 bp to +1 bp (Figure 3b). The most common indel was a 3 bp deletion present at 6% and there was some variation in where the DSB occurred. Some variation may be genuine, but for the most common mutations, DSB variance could be a sequencing artifact due to the start and end of the deletion both being a guanine nucleotide. *PDS2* had a 2.3% editing efficiency, with four variants (present at >0.1% of sequences) ranging from −4 bp to +1 bp (Figure 3d). The highest frequency indel was a 1 bp deletion present at 1.2%.

## Discussion

These results represent the first instance of DNA-free RNP-mediated genome editing in raspberry. The transfection efficiency of 19% with pre-assembled CRISPR RNPs is substantial and in line with existing research. A meta-analysis found that out of 36 published examples of protoplast transfection with PEG, the average transfection efficiency was 17% (see table 2 in ^44^. If gene edited protoplasts can be regenerated into raspberry plants, a 19% transfection efficiency would likely result in higher recovery rate of transformed plants than *Agrobacterium*-based methods in raspberry, which is currently between 0.5-2% ^28,45^. The development of novel protoplast-based transfection protocols in crop species has increased rapidly in recent years, with new methods in grapevine ^24,46,47^, tomato ^48,49^, strawberry ^17,50^, lettuce ^40^, and potato ^51^, reflecting widespread interest. As genome-editing research accelerates in *Arabidopsis*, tobacco, wheat and rice ^52^, it will be vital to invest in method development in a wider range of crops such as raspberry to ensure that NGBTs can be fully deployed.

This study demonstrates the amenability of raspberry to protoplast isolation, with slightly higher protoplast yields (average 5 × 10^6^ protoplast g^-1^) than previously reported (3.93 × 10^6^ g^-1^)^34^. Yields from BWP102 are comparable to recent reports in woody species such as grapevine (6.6 × 10^6^ g^-1^) ^24^ though lower than the 1 × 10^7^ protoplast g^-1^ seen in *Arabidopsis* ^16^. Adapting the method of Yoo et al. (2007) ^16^ to raspberry demonstrates the versatility of their protocol which has been fundamental in reigniting interest in protoplast research. The most important variable for successful protoplast isolation in raspberry, as found elsewhere ^16,17^, is the tissue quality used as a protoplast source. The outbreeding nature and high heterozygosity of raspberry adds complexity, compared to many other inbreeding species, as seeds cannot be used to grow sterile, soft, young plants for enzymolysis. As such, each raspberry seed is genetically different, and the genetic backgrounds of elite cultivars must be preserved to maintain berry consistency and quality. Therefore, vegetative stem cultures are a good alternative protoplast source. However, stem cultures are time and energy intensive, susceptible to contamination, and the health/quality of the canes is hugely influential on the protoplast yield. Although not systematically explored here, anecdotal evidence suggests indicators that improve protoplast yield include bright green stems that had deep red thorns and healthy lower leaves, alongside rapid growth of plantlets (within 10 days) from axillary buds in tissue culture. In future, isolation from raspberry callus could be an alternative protoplast source ^53^ to solve issues such as contamination and yield inconsistency.

Raspberry tissue remained mostly undigested despite 16 hrs of digestion, in contrast to *Arabidopsis* where leaves almost fully disintegrate into the enzymolysis solution ^23^. As a woody species, raspberry likely has higher concentrations of structural polymers, such as lignin and cellulose, compared to model species. We hypothesise that levels of structural polymers in raspberry have a positive correlation to age and abiotic stress ^54^. Therefore, high levels of structural polymers may interfere with protoplast isolation ^39^, resulting in lower yields from plantlets derived from older canes in poor health.

Pre-assembled RNPs were used in this study for several reasons. Primarily, non-transgenic crops are preferable as new legislation in England ^25^ permits only precision bred crops with mutations that could theoretically be produced by natural mutagenesis. Similar legislation is planned in the European Union ^26^. The genomes of genetically modified crops inevitably contain foreign DNA (*Agrobacterium* T-DNA, Cas9/gRNA DNA or plasmid sequences) which need to be removed through backcrossing. As previously stated, backcrossing is not practical in raspberry, and in many other soft fruits, as the species is highly heterozygous and outbreeding. Public perception of transgenic food also remains a major issue ^55^ and would likely influence marketability and eventual sales. Transgene-free cultivars generated by pre-assembled RNPs with enhanced phenotypes present a solution that research shows is more acceptable with consumers when used to improve sustainability or for societal benefit ^56^. Furthermore, the ability to rapidly introduce targeted non-transgenic mutations directly in clones with elite genetic backgrounds would be a significant advantage compared to the years of selection and trials required for conventional raspberry breeding. Floricane cultivars of raspberry take two years to fruit, thus RNP-based methods represent a great increase in breeding speed and accuracy. However, the above advantages still rely upon the elucidation of a protoplast regeneration protocol in raspberry ^34^.

Plasmid-derived RNPs would likely be effective in this protocol and would reduce costs long term. We have successfully tested plasmid YFP expression in BWP102 protoplasts (Figure S2). However, the risk of plasmid integration into the genome ^57^ and optimisation of Cas9/gRNA expression resulted in the preference of pre-assembled RNPs. Commercially synthesised Cas9 and gRNA enable reliable high purity synthesis and certification of sequence accuracy, which promotes high genome editing efficiency. Furthermore, it reduces the technical knowledge required to construct new plasmids for transfection. New gRNAs can be designed and synthesised rapidly if existing gRNAs prove ineffective. However, synthetic gRNAs and Cas9 are relatively expensive, therefore once gRNA designs are validated it may be more cost-efficient for any further work to synthesise and purify RNPs in-house.

The cleavage test validated the efficacy of the gRNAs *in vitro*. Screening of gRNAs through cleavage tests is recommended prior to protoplast transfection ^39^ to inexpensively validate the activity of gRNAs using minimal components. Availability of high-quality cultivar-specific genome assemblies were crucial for the design of highly specific gRNAs and oligonucleotides with no sequence mismatches or off-target complementarity; the assembly used in this study is intended for publication (Kevei et al., unpub.). Other raspberry cultivar genome sequences are available ^58,59^ which can be used for primer and gRNA design.

Quantifying genome editing efficiency in an amplicon pool is complex. T7EI and Sanger sequencing/deconvolution were used initially, which revealed positive results for *PDS1*, therefore amplicon sequencing was additionally conducted for more comprehensive analysis. Sanger sequencing followed by computational deconvolution offers a low-cost alternative when NGS is financially unfeasible. Many free-to-use programs exist, namely TIDE ^42^, ICE ^60^ and DECODR ^61^. While useful, discrepancies in editing estimations between different programs mean they cannot be singularly relied upon ^62^. Thus, if only using Sanger sequencing, editing estimates should be complemented with more substantial *in vitro* proof through the T7EI digestion assay. Taken together, these two analysis techniques can estimate what genotypes are present in the population and their relative frequencies at low cost. In this study, the T7EI assay demonstrated successful genome editing with RNP1, however did not validate the evidence of genome editing with RNP2 from TIDE, likely because editing was at a much lower frequency. The additional third band present in the T7EI *PDS1* sample could be explained by the endonuclease binding to a proportion of the DNA, reducing mobility through the agarose gel ^63^. Only amplicon sequencing was sensitive enough to detect the low editing efficiency of RNP2. Comparing the results of deconvolution and NGS reveals deconvolution to be a rough estimation of true indels and their frequencies. Many correct indels were detected, however their frequencies within the sequence pool were generally not accurate. Until recently, amplicon sequencing has been prohibitively expensive, thus methods such as T7EI and deconvolution were appealing. However, the decreasing cost of sequencing technologies, use of short-read (<500 bp) sequencing and the lack of high levels of accuracy with alternative methods, as identified here, indicates that amplicon NGS is the best choice for the detection of genome editing.

Genome editing and protoplast regeneration methods in raspberry could improve the sustainability of production through reduced waste, improved productivity, and enable better adaptation to climate change through increased flood, drought and disease resistance. One current concern in raspberry is the lack of vernalisation increasingly seen in some production regions due to warmer winters ^1^. *PDS* was chosen as a proof of concept as null mutants will be phenotypically identifiable early in plant development once protoplast regeneration is elucidated. In tandem with chromosome-level genome assemblies, this method enables rapid editing of any loci within the raspberry genome. However, as widely documented ^18,49^, protoplast regeneration of gene edited elite lines remains the greatest challenge to commercial implementation. This study aims to instigate further research into NGBTs in *Rubus* species by providing a straightforward, effective and reproducible protoplast isolation and DNA-free CRISPR genome editing protocol.

## Conclusion

Genome editing offers a promising route to advanced breeding in raspberry. This study describes fundamental species-specific methods that enable implementation and quantification of transgene-free edits in single-celled protoplast with considerable editing efficiency. Avenues have been opened for future work on gene expression and function. If protoplast regeneration methods can be elucidated in raspberry, genetic improvement of many raspberry traits will become feasible, with substantial benefits for the sustainability and efficiency of raspberry production.

## Acknowledgements

This research was funded by BerryWorld Plus™. We thank Dr Yaomin Cai (Earlham Institute) and Dr Mehmet Fatih Kara (University of York) for their advice on protoplast isolation and transfection and Dr Tomasz Kurowski (Cranfield University) for his work on the genome assembly.

## Author Contributions

R.C. conducted all experimental work, method development and wrote the manuscript. Z.K. and A.T. supervised the project and revised the manuscript. Z.K. and A.T. in collaboration with BerryWorld Plus™ conceived and wrote the project proposal.

