## Supplementary Information for "DNA-free CRISPR Genome Editing in Raspberry (*Rubus idaeus*) through RNP-mediated Protoplast Transfection and Comparison of Indel Analysis Techniques"

Figure S1: Screenshot of deconvolution result from TIDE of *PDS1* (a) and *PDS2* (b). Note increase in aberrant sequence signal after gRNA1 expected cut site. (c, d) screenshot of Sanger sequencing chromatogram in Geneious Prime for *PDS1* (c) and *PDS2* (d), showing decrease in nucleotide consensus downstream of highlighted gRNA cut site in *PDS1* only.

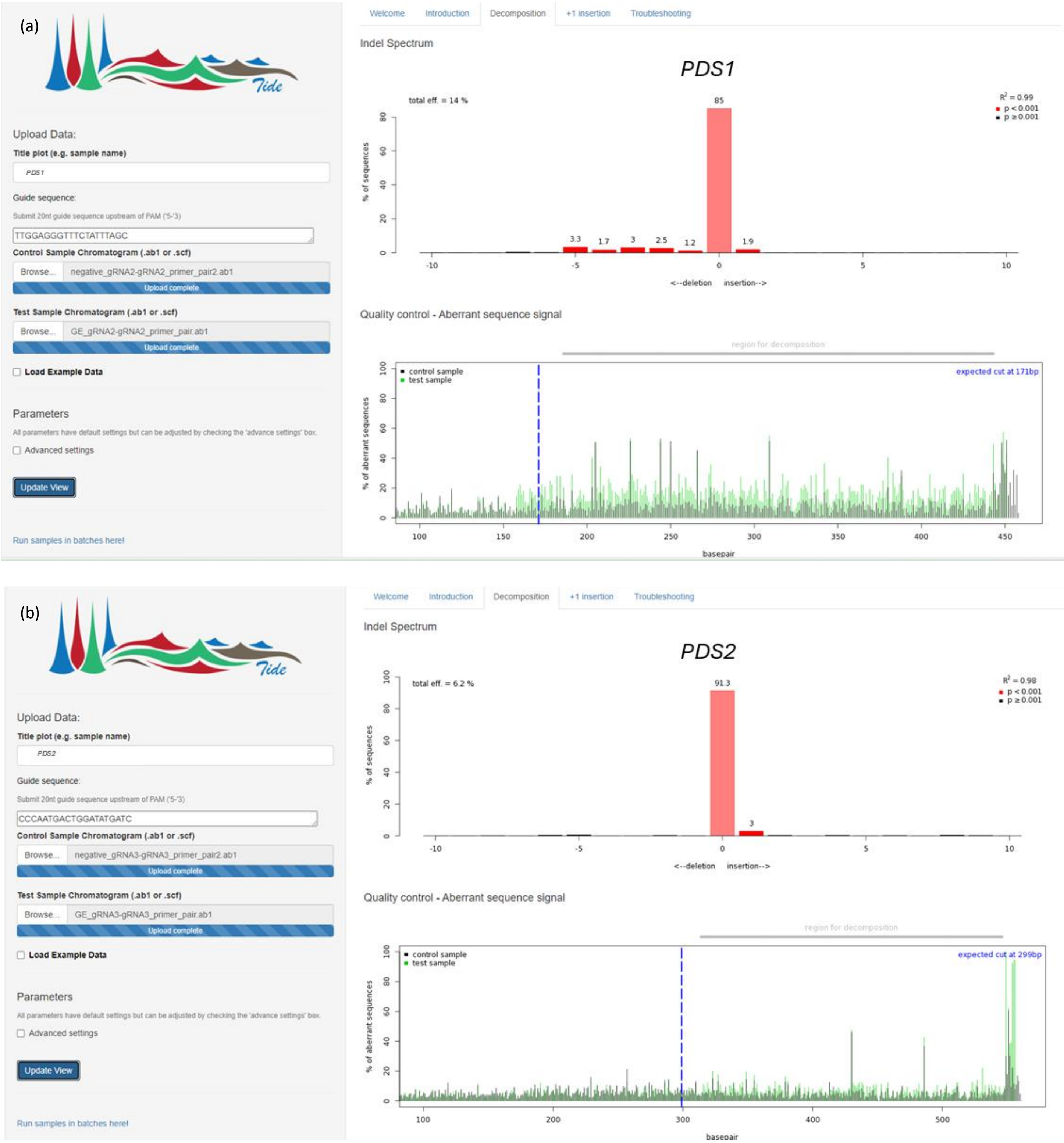

(c)

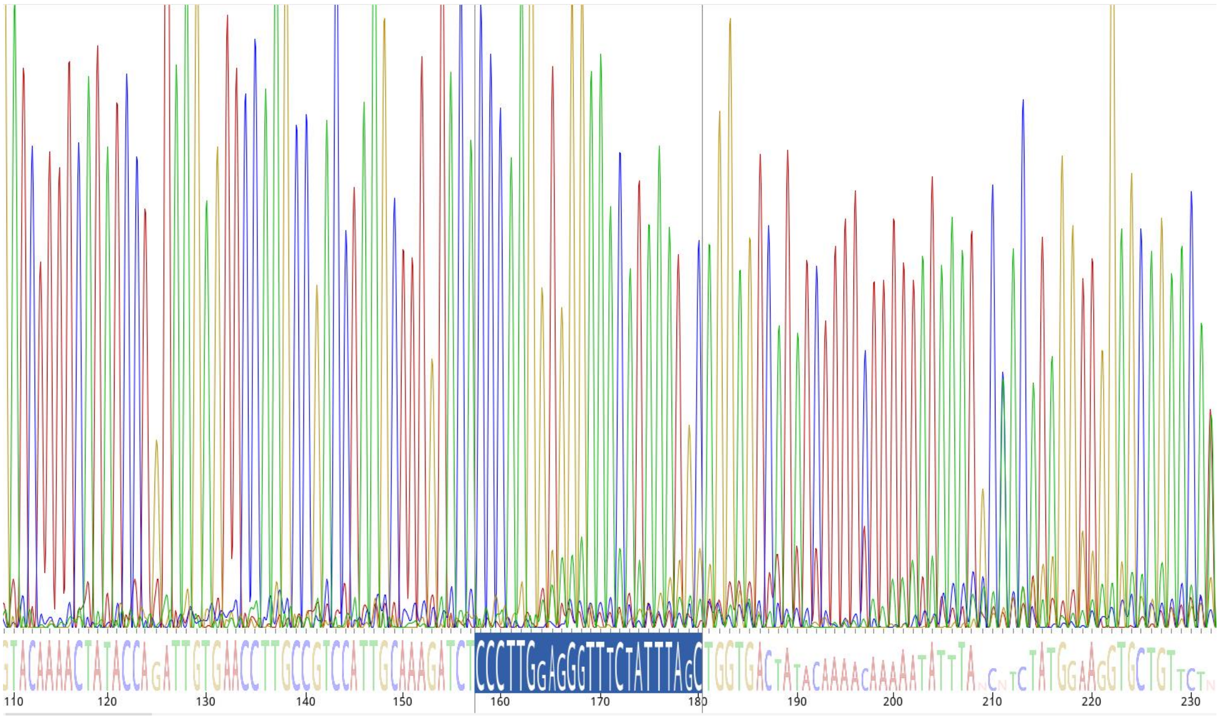

(d)

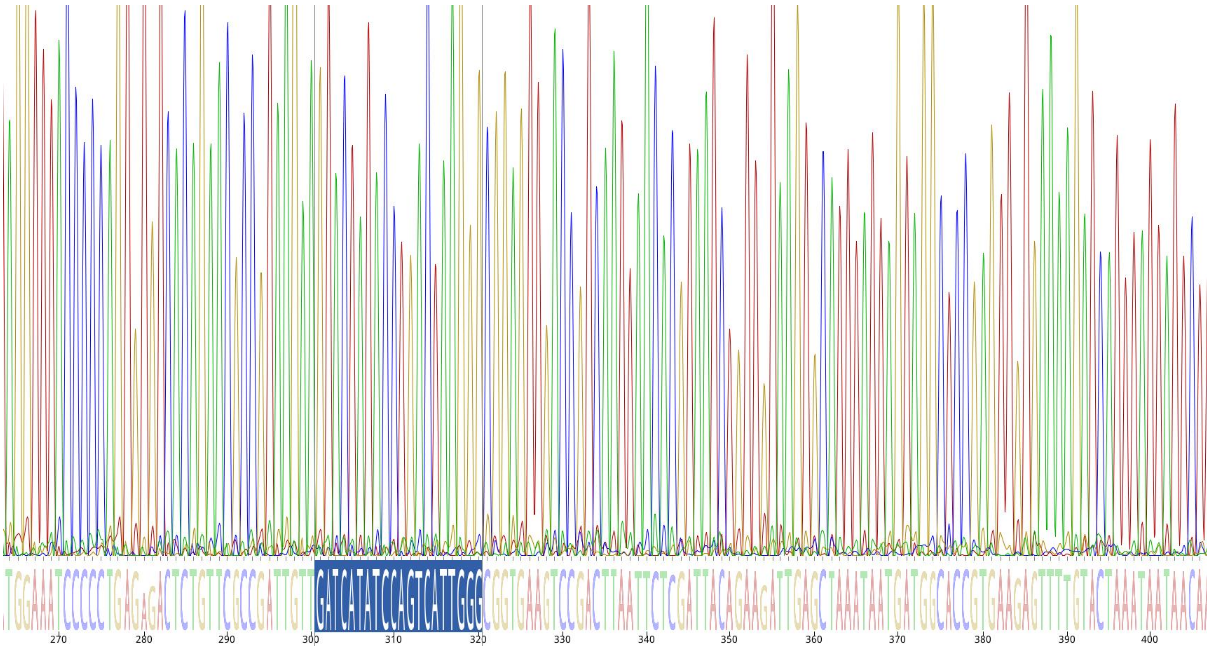

Figure S2: (a) Close up of BWP102 protoplast transfected with YFP plasmid; (b) YFP-containing plasmid used to transfect BWP102 protoplasts (c) 4x magnification image of sample of BWP102 protoplast showing widespread transfection. Error bar in (a) is 50µM and (c) is 200µM.

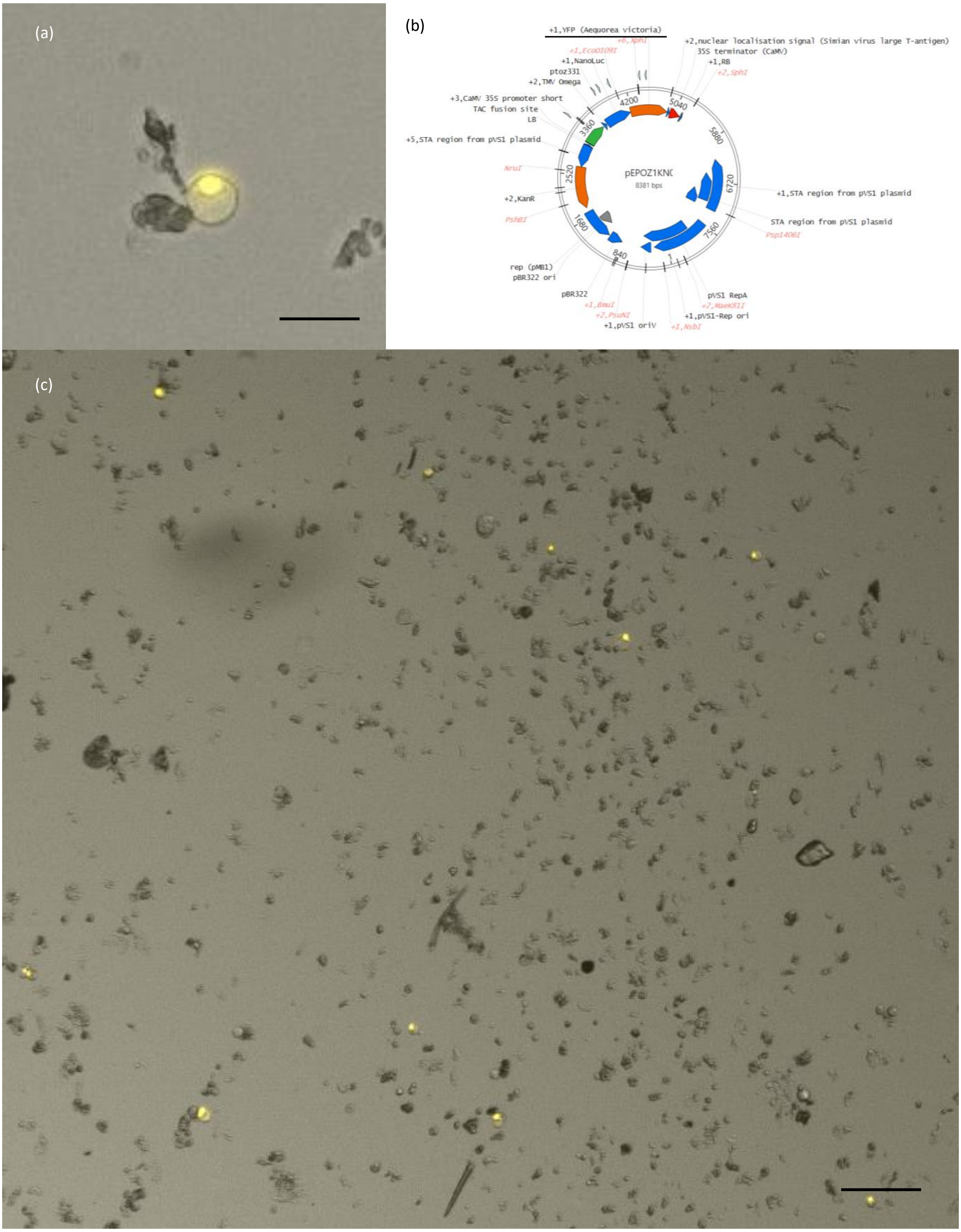
